# Complex synaptic and intrinsic interactions disrupt input/output functions in the hippocampus of *Scn1b* knockout mice

**DOI:** 10.1101/2023.04.29.538823

**Authors:** Jessica Hotard Chancey, Alisha A. Ahmed, Fernando Isaac Guillén, MacKenzie A. Howard

## Abstract

Mutations in the *SCN1B* gene have been linked to severe developmental epileptic encephalopathies including Dravet syndrome. *Scn1b* <u>k</u>nock<u>o</u>ut (KO) mice model *SCN1B* loss of function disorders, demonstrating seizures, developmental delays, and early death.*SCN1B* encodes the protein β1, an ion channel auxiliary subunit that also has roles in cell adhesion, neurite outgrowth, and gene expression. The goal of this project is to better understand of how loss of β1 alters information processing in the brain, resulting in seizures and associated cognitive dysfunction. Using slice electrophysiology in the CA1 region of the hippocampus from male and female *Scn1b* KO mice and <u>w</u>ild-type (WT) littermates, we found that processing of physiologically relevant patterned <u>S</u>chaffer <u>c</u>ollateral (SC) stimulation produces larger, prolonged depolarizations and increased spiking in KO neurons compared to WTs. KO neurons exhibit enhanced intrinsic excitability, firing more action potentials with current injection. Interestingly, SC stimulation produces smaller, more facilitating excitatory and inhibitory postsynaptic currents in KO pyramidal neurons, but larger postsynaptic potentials with the same stimulation. We also found reduced intrinsic firing of parvalbumin-expressing interneurons and disrupted recruitment of both parvalbumin- and somatostatin-expressing interneurons in response to patterned synaptic stimulation. Neuronal information processing relies on the interplay between synaptic properties, intrinsic properties that amplify or suppress incoming synaptic signals, and firing properties that produce cellular output. We found changes at each of these levels in *Scn1b* KO pyramidal neurons, resulting in fundamentally altered information processing in the hippocampus that likely contributes to the complex phenotypes of *SCN1B*-linked epileptic encephalopathies.

**Significance statement:** Genetic developmental epileptic encephalopathies have limited treatment options, in part due to our lack of understanding of how genetic changes result in dysfunction at the cellular and circuit levels. *SCN1B* is a gene linked to Dravet syndrome and other epileptic encephalopathies, and *Scn1b* knockout mice phenocopy the human disease, allowing us to study underlying neurophysiological changes. Here we found changes at all levels of neuronal information processing in brains lacking β1, including intrinsic excitability, synaptic properties, and synaptic integration, resulting in greatly enhanced input/output functions of the hippocampus. Our study shows that loss of β1 results in a complex array of cellular and network changes that fundamentally alters information processing in the hippocampus.

## Introduction

Neurons extract meaningful information from ongoing neural activity by integrating inputs from up to tens of thousands of synapses and transforming that information into appropriate outputs in the form of action potentials. The amplitude, kinetics, and summation of synaptic potentials is shaped by presynaptic release properties, and postsynaptic ion channels expressed in spine, dendritic, and somatic compartments. The shape and extent of the dendritic arbor, and the relative timing and location of excitatory and inhibitory inputs also affect how inputs interact (Spruston, 2008; Kole and Stuart, 2012; Stuart and Spruston, 2015). Thus, information processing by a neural circuit such as the hippocampus is a finely balanced interplay of many variables, and pathological disruption of these variables that alter input/output transformation can result in neurological disease.

Understanding the full etiology and uncovering therapeutic pathways for neurological diseases requires defining the causative insult and primary phenotypes, as well as secondary and tertiary effects, including how cellular and circuit processing is changed. The variety and complexity of epilepsy disorders illustrate the difficulty of this task. In addition to a wide variety of seizure phenotypes, epilepsy disorders present with a unique constellation of neuropsychiatric impairments. Symptomology is complex even when the causative insult seems simple in nature (e.g., a single point mutation). Many epileptic encephalopathies are monogenic disorders that cause drug-resistant seizures and a wide range of cognitive and developmental disabilities (Howard and Baraban, 2017).

Loss of function of the *SCN1B* gene results in a range of Early Infantile Epileptic Encephalopathy disorders, including Generalized Epilepsy with Febrile Seizures Plus (GEFS+) and Dravet syndrome (Scheffer et al., 2007; Patino et al., 2009; Ogiwara et al., 2012; Aeby et al., 2019). *SCN1B* encodes the protein Beta 1 (β1), which was first described as a sodium channel auxiliary subunit (Isom et al., 1992). β1 modulates the trafficking and physiology of many voltage-gated Na^+^ (Hull and Isom, 2018) and K^+^ (Marionneau et al., 2012; Nguyen et al., 2012) channels vital for action potential initiation and dendritic excitability.β1 is also necessary for developmental neuronal patterning, neurite outgrowth, and cell adhesion (Brackenbury et al., 2008; Patino et al., 2011; Brackenbury et al., 2013), and a soluble cleavage product translocates to the nucleus and regulates gene expression (Haworth et al., 2022). β1 is also essential for mature brain function, as deletion of β1 in adult mice leads to epilepsy and premature death (O’Malley et al., 2019).

Our goal was to understand how loss of β1 alters the input/output functions of the hippocampus using the *Scn1b* knockout (KO) mouse model, which phenocopies many aspects of *SCN1B* loss-of-function disorders, including febrile and spontaneous seizures, ataxia, altered cardiac physiology, developmental delays, and early death (Chen et al., 2004; Lopez-Santiago et al., 2007). We measured synaptic integration, synaptic physiology, and intrinsic physiology in CA1 pyramidal neurons and GABAergic interneurons. Rather than a single phenotype that intuitively explains neural dysfunction, we found a constellation of changes to every level of neuronal information processing in the hippocampus. CA1 pyramidal neurons were hyperresponsive to theta burst-patterned synaptic input. We tested hypotheses that this might be due to intrinsic hyperexcitability or increased/unbalanced synaptic strength. We found that while CA1 pyramidal neurons fired more action potentials with current injections, intrinsic physiology changes were complex and indicated the involvement of multiple ion channel types. Both excitatory and inhibitory synaptic currents were reduced in KO pyramidal cells, but were normally balanced. Finally, we tested the hypothesis that inhibition may be failing to effectively control activity in pyramidal neurons. These experiments revealed that parvalbumin-expressing interneurons demonstrated decreased intrinsic firing, and that both parvalbumin- and somatostatin-expressing interneuron subtypes exhibited drastically reduced recruitment in response to theta-burst patterned synaptic input. Thus, the physiological phenotypes associated with loss of *Scn1b* are distributed across cell types and functional domains, and subtle changes to interacting physiological mechanisms combine to alter how synaptic information is processed, likely resulting in the complex etiology of epileptic encephalopathies.

## Materials and Methods

### Animals

All procedures were approved by the Institutional Animal Care and Use Committee at the University of Texas at Austin. *Scn1b* knockout mice were a generous gift from Dr. Lori Isom at the University of Michigan (Chen et al., 2004). The colony was maintained by crossing *Scn1b*^+/-^ with C57Bl/6J mice (Jackson Labs, Strain #000664). To generate experimental mice, we bred *Scn1b*^+/-^ x *Scn1b*^+/-^. Male and female *Scn1b*^-/-^ (KO) and *Scn1b*^+/+^ (WT) littermates were used for experiments at postnatal day (P) 15-20. To generate interneuron reporter mice, we crossed *Scn1b* knockout mice with PV-Cre (Jackson Labs, Strain # 17320) or SST-Cre (Jackson Labs, Strain #28864) and the Ai14-TdTomato reporter (Jackson Labs, Strain #007914). Pups were toe-clipped at P6-8 for identification, and tail tissue samples were taken for genotyping by PCR as in Chen et al. (2004), or were genotyped by TransnetYX, Inc. As previously reported,*Scn1b*^-/-^ mice in our colony had spontaneous seizures and stunted growth beginning around P10 and exhibited 100% mortality by P22 (Chen et al., 2004; Hull et al., 2020).

### Acute slice preparation

Mice were deeply anesthetized with a mix of ketamine (90 mg/kg; Acor) and xylazine (10 mg/kg; Dechra) intraperitoneally. Mice were transcardially perfused with ice-cold, oxygenated cutting solution, containing (in mM): 205 sucrose; 25 sodium bicarbonate; 2.5 KCl; 1.25 sodium phosphate; 7 MgCl_2_; 7 D-glucose; 3 sodium pyruvate; 1.3 ascorbic acid; 0.5 CaCl_2_. Brains were removed, hemisected, and the dorsal portion of the brain was removed using a scalpel blade, angled at ∼20°. The dorsal cut side was then mounted in the slicing chamber of a Leica VT1200 vibratome using superglue, and submerged in ice-cold, oxygenated cutting solution. 250 μm oblique-horizontal hippocampal slices were made. Slices incubated for 30 minutes at 37° C in holding solution containing (in mM): 125 NaCl; 25 sodium bicarbonate; 2.5 KCl; 1.25 sodium phosphate; 12.5 D-glucose; 2 MgCl_2_; 2 CaCl_2_; 1.3 ascorbic acid; 3 sodium pyruvate. Slices were then stored for at least 30 minutes, and up to 8 hours, in holding solution at room temperature before being used for recordings. Reagents for electrophysiology solutions were purchased from Sigma Aldrich.

### Whole-cell electrophysiology

Slices were transferred to the recording chamber and were continuously bathed in warm (32.5° C), oxygenated artificial cerebrospinal fluid (ACSF), containing (in mM): 125 NaCl, 25 NaHCO_3_, 2.5 KCl, 1.25 NaH_2_PO_4_, 1 MgCl_2_, 2 CaCl_2_. Only slices from the middle of the dorsal-ventral axis of the hippocampus were used for recordings. Patch pipettes of 3-6 MΩ were pulled from thin-walled borosilicate filamented glass (Sutter BF150-110-10) using a P-1000 puller (Sutter). For current clamp experiments, pipettes were filled with a potassium gluconate solution containing (in mM): 120 K-gluconate, 20 KCl, 10 HEPES, 4 NaCl, 1 EGTA; 4 Mg-ATP, 0.3 Na-GTP, 7 phosphocreatine disodium salt hydrate. For voltage clamp experiments, the internal solution contained (in mM): 120 cesium methansulfonate; 4 NaCl; 6 CsCl; 10 HEPES; 1 EGTA; 4 Mg-ATP, 0.3 Na-GTP, 7 phosphocreatine disodium salt hydrate; 5 QX-314. 0.3% biocytin was included in the internal solution of some experiments for post-hoc recovery of recorded neurons. Slices were visualized using a Zeiss Axio Examiner with Dodt contrast optics and epifluorescence, and a Zeiss Axiocam 503 infrared digital camera. Data were acquired with a MultiClamp 700B amplifier (Molecular Devices) and an InstruTECH LIH8+8 (HEKA) digitizer at a sampling rate of 10 kHz, filtered at 4 kHz (Bessel). Pipette capacitance neutralization and bridge balance were applied, and liquid junction potential was not corrected for. Data were acquired using Axograph software. After measuring resting membrane potential in current clamp experiments, a holding current was applied to give all neurons a baseline membrane potential of ∼-70 mV. We excluded cells with a series resistance > 25 MΩ or if the series resistance changed by more than 20% throughout the recording. Synaptic stimulation was delivered by a tungsten bipolar stimulating electrode (WE3ST30.1ASS, Microprobes) placed in the stratum radiatum near the CA2/CA1 border to stimulate the Schaffer collateral axons, and a constant current stimulus isolator (World Precision Instruments). Gabazine was purchased from Sigma Aldrich or Hello Bio. ZD7288 was purchased from Tocris or Hello Bio.

### Morphology

Slices containing biocytin filled neurons were fixed in 4% PFA (Electron Microscopy Science) overnight at 4°, and then stored in phosphate-buffered saline (PBS) until processed. Filled neurons were stained using Vectastain ABC-HRP and DAB kits (Vector Labs) and reconstructed using the Neurolucida Tracing Program (MBF Bioscience). Sholl analyses, using a 15 μm radius, were used to measure dendritic length and dendritic branching patterns. Neurons in which major dendritic branches had been cut were excluded from these analyses.

### Data analysis and statistics

Electrophysiology data were analyzed using Axograph software. Input resistance was measured as the slope of the linear portion of current-voltage plot using negative and subthreshold positive current steps. Spike threshold was defined as the membrane potential where dV/dt reached 10% of the maximum dV/dt. Action potential amplitude was taken as the difference between threshold and max voltage. Capacitance was calculated by measuring the time constant (τ) of repolarization from a −50 pA current step and dividing that value by the membrane resistance. Data were analyzed using either Student’s t-tests, Mann-Whitney tests, or analysis of variance (ANOVA) followed by post-hoc multiple comparisons tests, as appropriate using GraphPad Prism 9. N values used for statistics are number of cells, but number of animals is also listed for each experiment. No more than three cells from a single animal were used for any experiment. All data are presented as mean ± SEM.

## Results

A neuron’s fundamental role is to receive incoming synaptic inputs, properly filter and integrate those inputs, and produce an appropriate output in the form of action potentials. To examine basic cellular information processing in the hippocampus of *Scn1b* KO mice we performed a synaptic integration study, in which theta burst stimulation (5 pulses at 100 Hz per burst, repeated 5 times at 5 Hz; hash marks below example traces in Figure 1) was delivered to <u>S</u>chaffer <u>c</u>ollateral (SC) axons and the resulting postsynaptic voltage response was measured in CA1 pyramidal neurons. Stimulation intensity was normalized to the lowest stimulation that gave a reliable postsynaptic response of 1.5-2 mV for a single stimulation (1x, Figure 1A, left; inset shows an expanded view of the first burst). Using that minimal stimulation intensity, we found that *Scn1b* KO neurons showed enhanced temporal synaptic integration compared to WT littermates, measured as the area under the curve of the voltage response with spikes truncated (Figure 1B left; main effect of genotype: F_(1, 20)_ = 6.876, p = 0.016; main effect of theta cycle: F_(1.3, 26.4)_ = 3.796, p = 0.052, interaction: F_(4,80)_ = 0.222, p = 0.925; two-way <u>r</u>epeated <u>m</u>easures (RM) ANOVA). As we increased the stimulation intensity by multiples of the minimal intensity, the response in KO neurons drastically increased in amplitude (Figure 1B; middle is 3 times minimal stim: main effect of genotype: F_(1, 20)_ = 19.3, p = 0.0003; theta cycle: F_(1.6, 32)_ = 2.93, p = 0.079, genotype x theta cycle: F_(4,80)_ = 0.605, p = 0.660; right is 5x minimal stim: main effect of genotype: F_(1, 20)_ = 12.89, p = 0.002; theta cycle: F_(1.6, 33)_ = 1.894, p = 0.172, genotype x theta cycle: F_(4,80)_ = 0.730, p = 0.574). The probability of firing and the number of action potentials also increased with stim intensity in KO neurons, whereas, this range of stimulation did not drive action potential firing in WT neurons (Figure 1C: main effect of genotype: F_(1,20)_ = 25.3, p <0.0001; main effect of stim intensity: F_(2.8, 56.17)_ = 7.576, p = 0.0003; intensity x genotype: F_(4, 80)_ = 7.56, p < 0.0001; Figure 1D: main effect of genotype: F_(1,20)_ = 3.740, p = 0.067; main effect of stim intensity: F_(1.8, 35.8)_ = 3.189, p = 0.058; intensity x genotype: F_(4, 80)_ = 3.189, p = 0.018; two-way RM ANOVA). This indicates a major shift in input/output processing in CA1 pyramidal neurons lacking *Scn1b*, which intuitively relates to the seizures and changes in cognitive function exhibited by people with *SCN1B*-linked encephalopathies.

**Figure 1.**
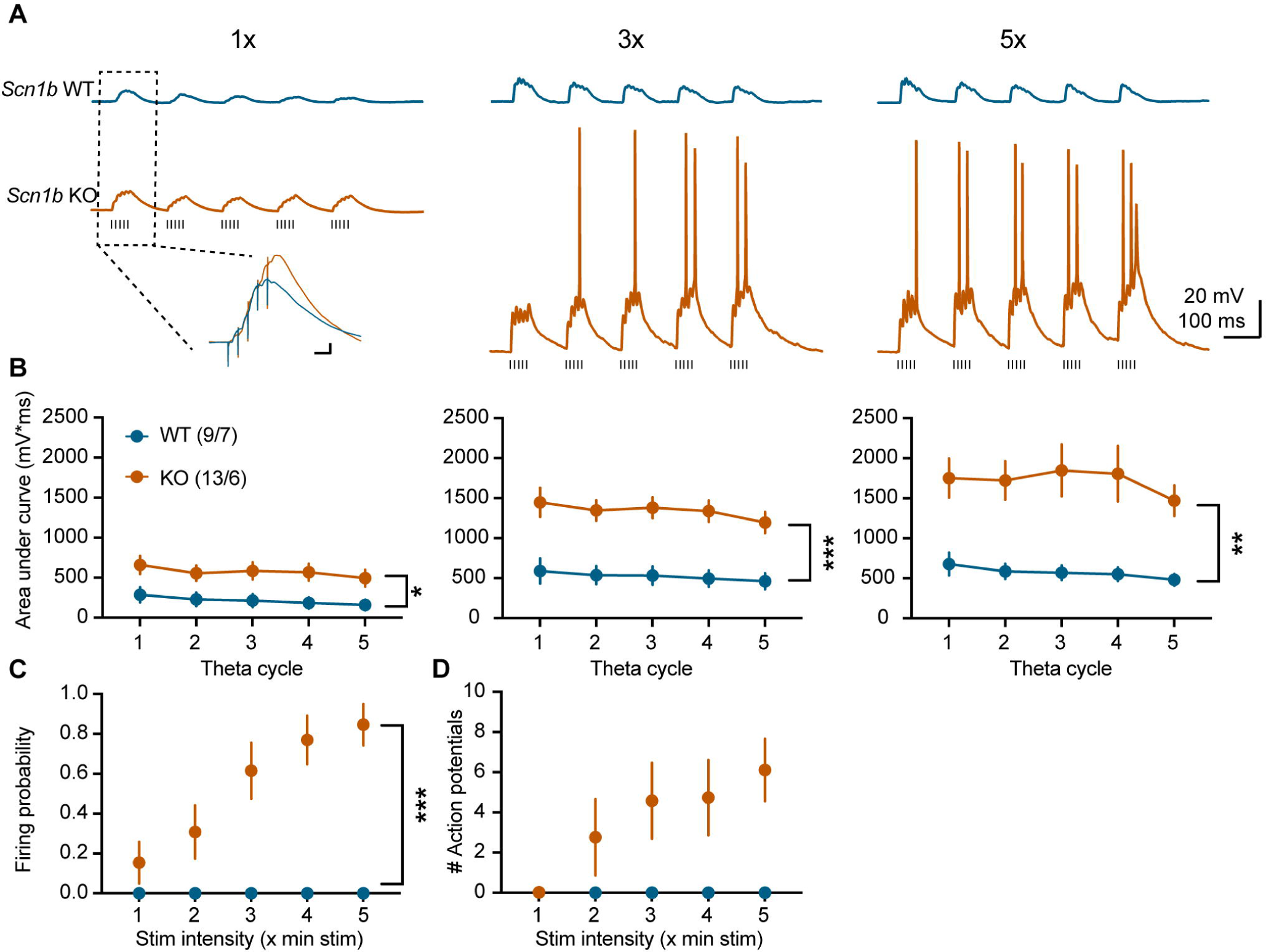
Aberrant temporal summation in *Scn1b* KO CA1 pyramidal neurons. **A)** Postsynaptic potentials measured in CA1 pyramidal neurons in response to theta burst stimulation of Schaffer collateral axons (5 pulses at 100 Hz, 5 bursts, 200 ms inter-burst interval; designated by hash marks below traces) from *Scn1b* WT (blue, top) and KO (orange, bottom) neurons. Stim artifacts were removed for clarity. The stimulation intensity was normalized such that the first stimulation of the first burst was 1.5-2 mV. Inset shows expanded view of first theta burst with stim artifacts to demonstrate that the first pulse was the same amplitude across experiments (inset scale = 2 mV, 20 ms). Stimulation intensity was increased by multiples of the minimal stimulus to 3x (middle) and 5x (right). **B)** At minimal stim intensity (1x; left) *Scn1b* KO neurons had larger responses, i.e., stronger synaptic integration, compared to WT, measured as area under the curve with spikes truncated. As the stimulation intensity was increased, the enhanced integration became more prominent (middle = 3x, right = 5x minimal stim).**C)** The firing probability and **(D)** number of action potentials (including 0s) increased with increasing stimulation intensity in KO neurons, whereas WT neurons did not fire with this stimulation protocol. **B-D)** N = (cells/mice). * p < 0.05; ** p < 0.01; *** p < 0.001, main effects of genotype; two-way RM ANOVAs.

We next tested if intrinsic firing properties may underly the increased firing shown in Figure 1. Current steps from 0-300 pA, in 25 pA increments, 250 ms long, were injected into CA1 pyramidal neurons and firing properties were measured. Indeed, *Scn1b* KO pyramidal neurons fired more action potentials with increasing current injections compared to WT, but only with larger current steps (Figure 2 A-B; main effect of current step: F_(1.6, 45.7)_ = 115.0, p < 0.0001; genotype: F_(1, 29)_ = 4.736, p = 0.037; current x genotype: F_(12, 348)_ = 3.485, p < 0.0001; two-way RM ANOVA with post-hoc Sidak’s multiple comparisons test). We measured the rheobase (minimum amount of current needed to drive firing) in the cells from Figure 2B and in a second set of cells which were given current steps in 5 pA increments to more accurately assess the rheobase (n = 10 WT; 12 KO). These data are combined in Figures 2C-E because there were no significant differences between experiments. There was no difference in rheobase between genotypes (Figure 2C; WT: 130.2 ± 12.5 pA, n = 23 cells from 10 mice; KO: 132.3± 9.4 pA, n = 31 cells from 16 mice; t_(52)_ = 0.133, p = 0.894). There was also no difference in threshold voltage of the first action potential (Figure 2D; WT: −37.3 ± 0.9 mV; KO: −36.8 ± 0.7 mV; t_(52)_ = 0.425, p = 0.673) or resting membrane potential between genotypes (Figure 2E; WT: −61.3± 1.3 mV; KO: −61.5 ± 0.9 mV; t_(52)_ = 0.158, p = 0.875). However, the amplitude of the first action potential (measured at rheobase) was smaller (Figure 2F; WT: 80.33 ± 2.22 mV; KO: 73.72 ± 1.30 mV; t_(52)_ = 2.715, p = 0.009) and wider, measured as the width at half of the maximum amplitude, in KO pyramidal cells compared to WT (Figure 2G; WT: 1.62 ± 0.10 ms; KO: 2.02 ± 0.12 ms; t_(52)_ = 2.528, p = 0.015).

**Figure 2.**
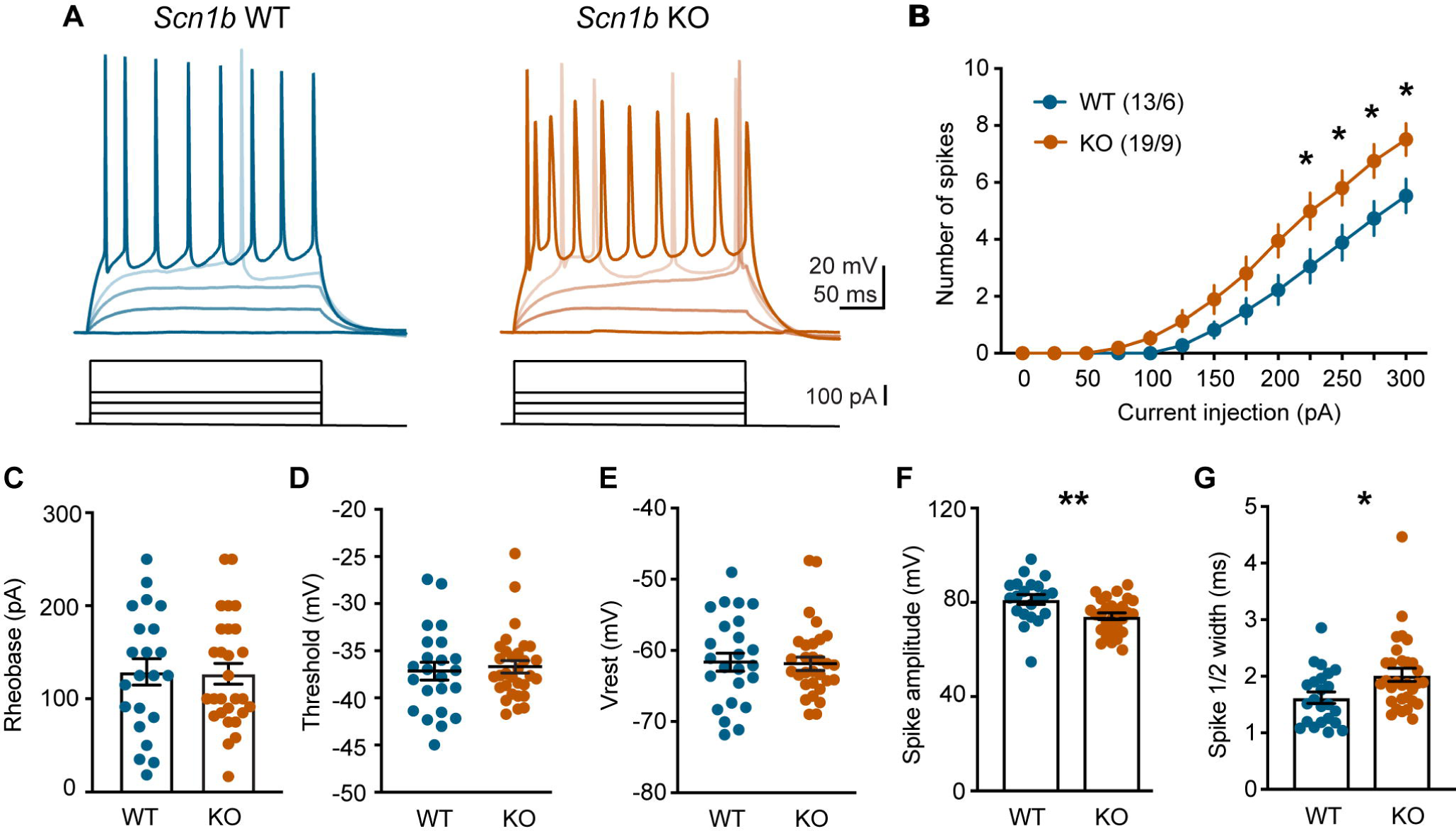
Increased action potential firing in *Scn1b* KO CA1 pyramidal neurons. **A)** Example voltage responses to increasing current injections from *Scn1b* WT (left, blue) and KO (right, orange) pyramidal cells. **B)** KO neurons fired more with increasing current injections compared to WT neurons. N = cells/mice. *p < 0.05 Sidak’s multiple comparisons test. **C)** KO CA1 pyramidal neurons did not exhibit changes to rheobase, **(D)** threshold voltage, or **(E)** resting membrane potential (Vrest). **F)** The first action potential (measured at the rheobase current step) had a lower amplitude and **(G)** wider half-width in KO neurons compared to WT. *p < 0.05; **p < 0.01; t-test.

We next investigated if the increase in firing of CA1 pyramidal neurons could be attributed to membrane properties. Using a series of negative current steps we measured passive membrane properties in pyramidal neurons (Figure 3A) and found that at resting membrane potential there was no difference in membrane resistance between WT and KO neurons (WT: 128.0 ± 19.3 MΩ, n = 13; KO: 137.4 ± 11.3 MΩ, n = 19; t_(30)_ = 0.449, p = 0.657, unpaired t-test; Figure 3A). The membrane capacitance was lower in KO neurons compared to WT (WT: 159.1 ± 12.0 pF; KO: 123.4 ± 6.8 pF; t_(30)_ = 2.775, p = 0.009, unpaired t-test; Figure 3B), suggesting that KO neurons may be smaller than WT neurons. Another observation from the negative current steps was that the sag voltage was larger in KO neurons compared to WT (Figure 3C-D; sag amplitude measured at the smallest hyperpolarizing step in which the neuron reached −90 mV; WT: 4.56 ± 0.47 mV; KO: 5.64 ± 0.19 mV; t_(30)_ = 2.414, p = 0.022). An increase in voltage sag is an indicator of increased <u>h</u>yperpolarization activated and <u>c</u>yclic-<u>n</u>ucleotide-gated (HCN) channel activity. HCN channels are cation channels that open at hyperpolarized potentials, and therefore, decrease membrane resistance and depolarize the membrane. Increasing HCN activity has been shown to reduce excitability and membrane resistance of pyramidal neurons (Poolos et al., 2002; Fan et al., 2005), whereas downregulation of HCN channels has been linked to hyperexcitability and increased membrane resistance in several epilepsy models (Chen et al., 2001; Strauss et al., 2004; Jung et al., 2007). Because there was no difference in membrane resistance (Figure 3A), but lower capacitance (Figure 3B), we hypothesized that *Scn1b* KO CA1 pyramidal neurons may be smaller, which typically correlates with higher membrane resistance, but that increased somatic HCN channels in KO neurons normalizes the membrane resistance. Indeed, when HCN channels were blocked with the antagonist ZD7288 (20 μM), membrane resistance increased (Figure 3 E, F; main effect of drug: F_(1,29)_ = 11.06, p = 0.002), and the increase was larger in KO neurons compared to WT (genotype x drug: F(_1, 29)_ = 4.277, p = 0.048); genotype: F_(1,29)_ = 2.007, p = 0.167; two-way RM ANOVA with post-hoc Sidak’s multiple comparisons test). ZD7288 significantly increased the resistance of KO neurons (p = 0.032, Sidak’s multiple comparisons test), but not WT neurons (p = 0.994), consistent with previous data showing that wild-type CA1 pyramidal cells have few HCN channels at the soma (Magee, 1998). Sag voltages were larger in KO neurons, but ZD7288 similarly blocked sag in both genotypes (Figure 4E; WT: 87.37 ± 3.12% block; KO: 74.81 ± 6.09%; p = 0.545, Mann-Whitney test; summary data not shown). To further test this hypothesis, we measured membrane resistance while varying the membrane potential, and found an increased slope in the KO data when resistance is plotted against membrane potential (Figure 3 G-H; main effect of voltage: F_(1.77, 35.4)_ = 121.1, p < 0.0001; genotype: F_(1,20)_ = 0.292, p = 0.595; voltage x genotype: F_(4,80)_ = 4.569, p = 0.02; two-way RM ANOVA), suggesting that membrane resistance increases more in KO neurons as the cell depolarizes and HCN channels deactivate. These data together suggest that *Scn1b* KO CA1 pyramidal neurons likely have more compact somata than WT, but somatic HCN channel number or activity is increased, normalizing the input resistance and rheobase. However, at more depolarized potentials, where HCN channels are deactivated, KO neurons exhibit increased excitability (Figure 2B, current steps > 200 pA). Overall, our intrinsic physiology data indicate modest hyperexcitability in these neurons, but the mixed changes in somatic membrane properties, lack of change to rheobase, threshold, or action potential number in response to smaller suprathreshold steps were surprising given the large increases in the input/output function to synaptic stimulation.

**Figure 3.**
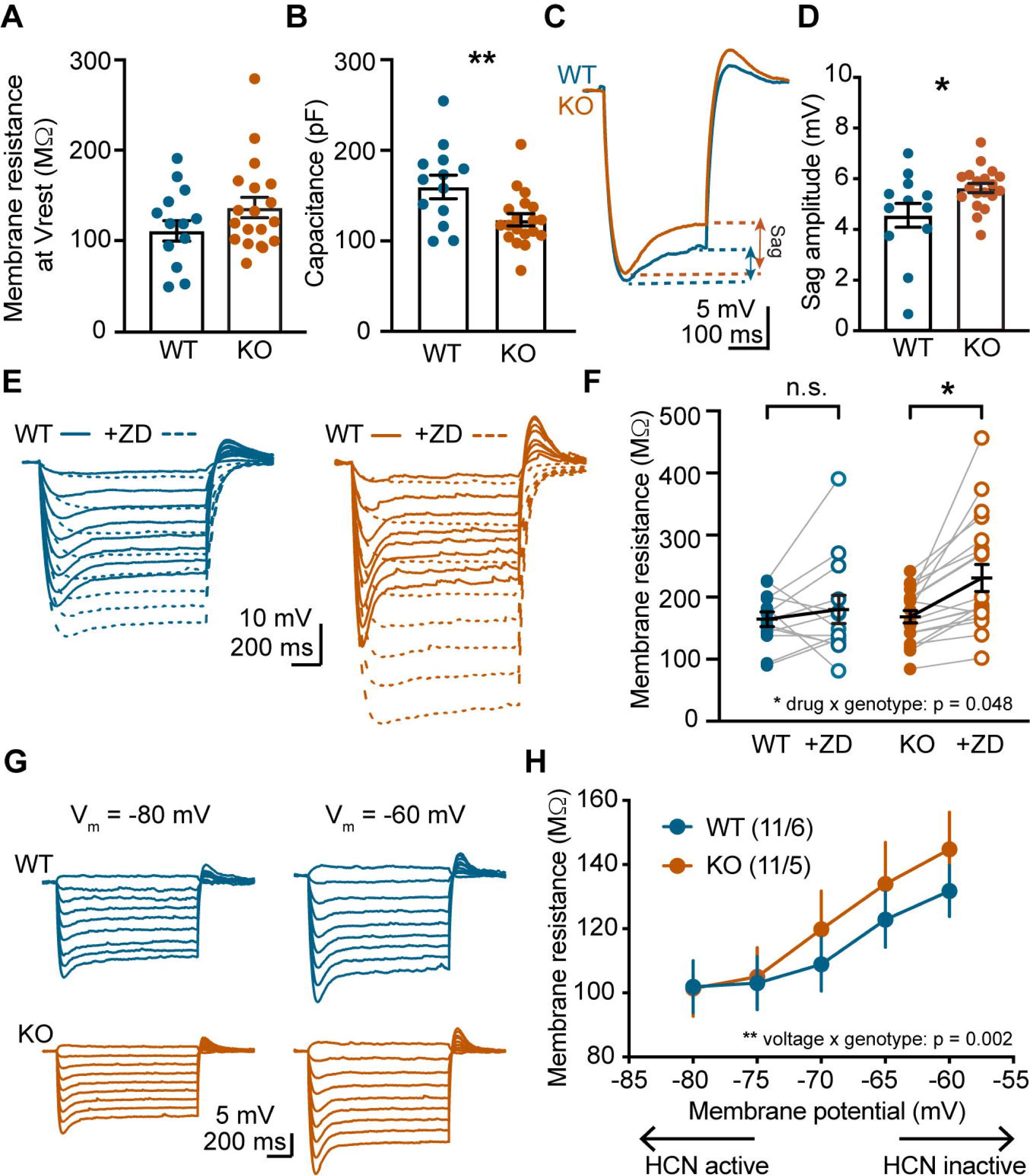
Increased HCN activity normalizes membrane resistance in *Scn1b* KO pyramidal neurons. **A)** Membrane resistance was not different, while capacitance **(B)** was decreased in KO pyramidal cells compared to WT at resting membrane potential. ** p < 0.01, t-test.**C)** Example voltage traces of a hyperpolarizing current step to −90 mV from a WT (blue) and KO (orange) pyramidal cell. Sag amplitude = peak negative voltage – steady state voltage. **D)** KO neurons show greater mean sag amplitude to current steps that cause peak hyperpolarization to −90 mV. * p < 0.05, t-test. **E)** Example voltage responses to hyperpolarizing current steps (−10 to −150 pA, 20 pA increments, 1 s) from WT (left, blue) and KO (right, orange) pyramidal cells in the absence (solid lines) and presence of 20 µM ZD7288 (dashed lines). **F)** Blockade of HCN channels with ZD7288 has a larger effect on membrane resistance of KO neurons compared to WT. N = 12 cells from 5 mice WT, 18 cells from 10 mice KO. n.s. = not significant (p > 0.05); * p < 0.05, Sidak’s multiple comparisons test. **G)** Example voltage traces in response to currents steps (+10 to −150 pA, 20 pA increments, 1 s) while holding the cell at varying membrane potentials (left = −80 mV; right = −60 mV). **H)** Summary data of membrane resistance vs. membrane potential. N = (cells/mice).

**Figure 4.**
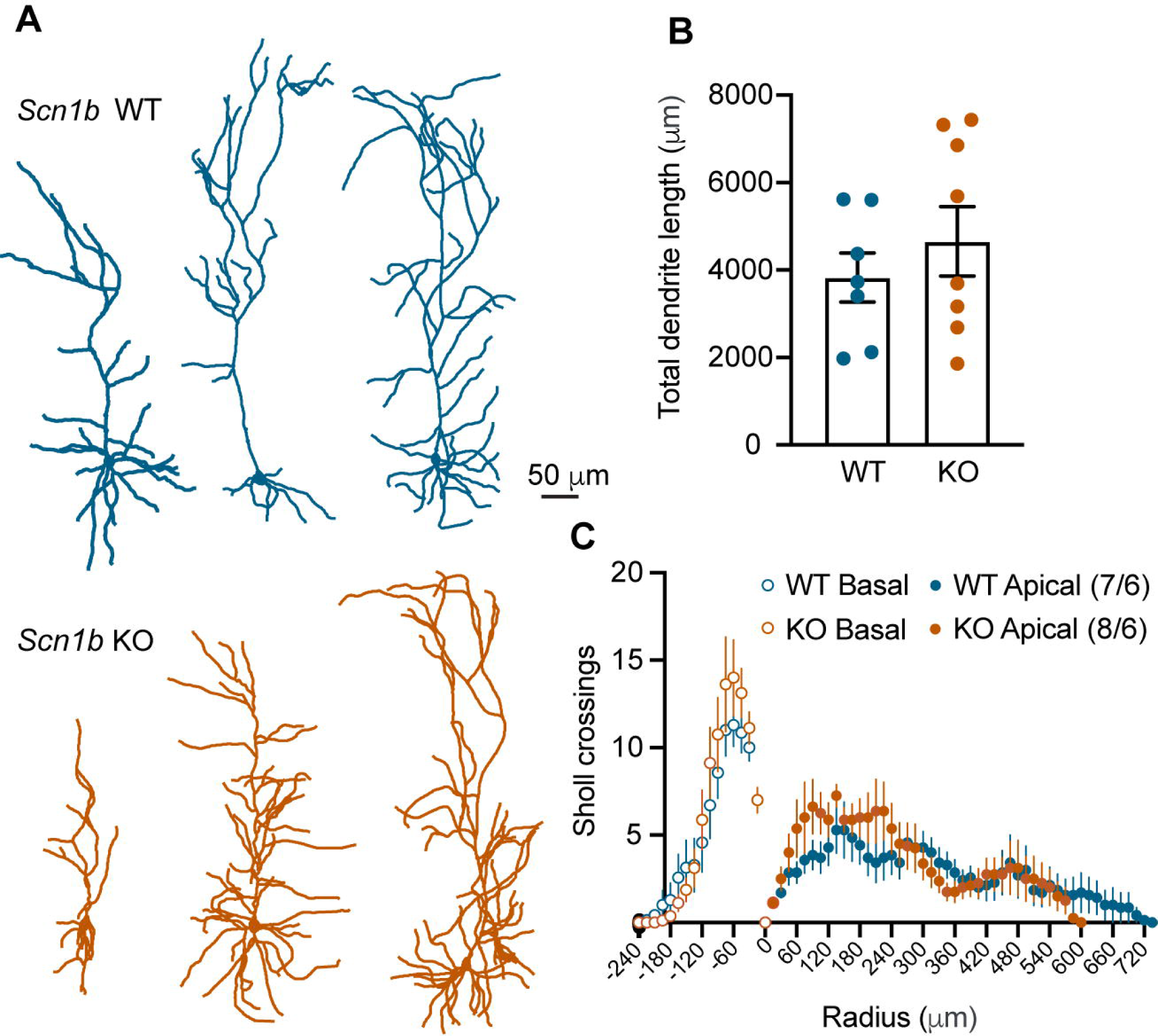
No change in dendritic structure in KO pyramidal neurons. **A)** Example reconstructions of *Scn1b* WT (top, blue) and KO (bottom, orange) CA1 pyramidal neurons. **B)** No difference in total dendrite length between genotypes. **C)** No difference in number of Sholl crossings using 15 μm radii for apical (solid circles) or basal dendrites (open circles). N = (cells/mice).

A previous study found that increased excitability was associated with reduced dendritic arborization in mutant *Scn1b* subicular pyramidal neurons (Reid et al., 2014). Furthermore, *Scn1b* KO mice have deficits in neurite outgrowth (Brackenbury et al., 2008). As such, we hypothesized that the decreased capacitance and hyperexcitability in *Scn1b* knockout CA1 pyramidal neurons could be due in part to reduced dendritic arborization. Recorded neurons were filled with biocytin and reconstructed. CA1 pyramidal cell dendritic structures are highly variable at this early age, as revealed by WT and KO neuron reconstructions (Figure 4A). There was no difference in total dendritic length (Figure 4B; t_(13)_ = 0.828, p = 0.423, t-test), or number of crossings as a function of distance from the soma between genotypes (Figure 4C; main effect of Sholl radius: F_(65, 849)_ = 15.83, p < 0.0001; main effect of genotype: F_(1, 849)_ = 3.415, p = 0.065; genotype x radius: F_(65, 849)_ = 0.774, p = 0.904; two-way ANOVA). These data indicate that the enhancement in excitability and synaptic integration were not associated with gross changes in dendritic morphology.

The subtle differences in KO intrinsic excitability described above are unlikely to fully account for the dramatic increase in synaptic integration seen in Figure 1. We next tested the hypothesis that excitatory synaptic inputs to *Scn1b* KO neurons were enhanced, or the balance of excitation and inhibition was altered. First, we used voltage clamp to isolate <u>e</u>xcitatory <u>p</u>ost<u>s</u>ynaptic <u>c</u>urrents (EPSCs) by holding the membrane at the reversal potential for GABA (−70 mV). We stimulated the SCs, normalizing to the lowest stimulation intensity that reliably evoked a postsynaptic current (∼40 pA EPSC in both genotypes), and then increased the intensity by multiples of that threshold to generate an input/output curve. Counter to expectations following current clamp experiments, the growth curve of the EPSC input/output curve was significantly lower in *Scn1b* KO neurons compared to WT (Figure 5 A-B; main effect of stim intensity: F_(8, 216)_ = 36.38, p < 0.0001; main effect of genotype: F_(1, 27)_ = 6.082, p = 0.020; stim intensity x genotype: F_(8,216)_ = 4.382, p < 0.0001; two-way RM ANOVA with post-hoc Sidak’s multiple comparisons test). The same neurons were then clamped at +10 mV to isolate inhibitory <u>p</u>ost<u>s</u>ynaptic <u>c</u>urrents (IPSCs), and using the same stimulation intensities as in Figure 5B, we found that the IPSCs were also substantially smaller in KO neurons compared to WT (Figure 2A, C; main effect of stim intensity: F_(8, 216)_ = 46.32, p < 0.0001; main effect of genotype: F_(1, 27)_ = 8.648, p = 0.007; stim intensity x genotype: F_(8,216)_ = 4.226, p < 0.0001; two-way RM ANOVA with post-hoc Sidak’s multiple comparisons test). There was no change in PSC kinetics between genotypes (EPSC rise: WT = 3.323 ± 0.327 ms, KO = 3.352 ± 0.403 ms, t_(31)_ = 0.038, p = 0.970; EPSC decay τ: WT = 9.867 ± 0.600 ms, KO = 8.140 ± 1.106 ms, t_(31)_ = 1.435, p = 0.161; IPSC rise: WT = 4.032 ± 0.491 ms, KO = 4.218 ± 0.627 ms, t_(31)_ = 0.238, p = 0.814; IPSC decay τ: WT = 27.700 ± 2.202 ms, KO = 19.390 ± 3.287 ms, t_(31)_ = 1.382, p = 0.177; t-tests). While both EPSCs and IPSCs were smaller in KO neurons, they were also more facilitating, measured as an increase in paired-pulse ratio (Figure 5A, D; 50 ms inter-stimulus interval at 3x threshold stim; EPSCs: WT = 1.052 ± 0.066, KO = 1.599 ± 0.281, p = 0.046; IPSCs: WT = 0.803 ± 0.102, KO = 1.699 ± 0.471, p = 0.0008, Mann-Whitney tests). EPSCs and IPSCs scaled down in a similar manner, as there was no difference in excitatory to inhibitory ratio between genotypes (Figure 5E; main effect of genotype: F_(1,27)_ = 3.371, p = 0.077, main effect of stim intensity: F_(8, 216)_ = 3.613, p = 0.0006; stim intensity x genotype: F_(8, 216)_ = 1.297, p = 0.246; two-way RM ANOVA). Finding significantly smaller PSCs and a lack of difference in excitatory to inhibitory ratio was unexpected given the significantly larger temporal synaptic integration seen in Figure 1.

**Figure 5.**
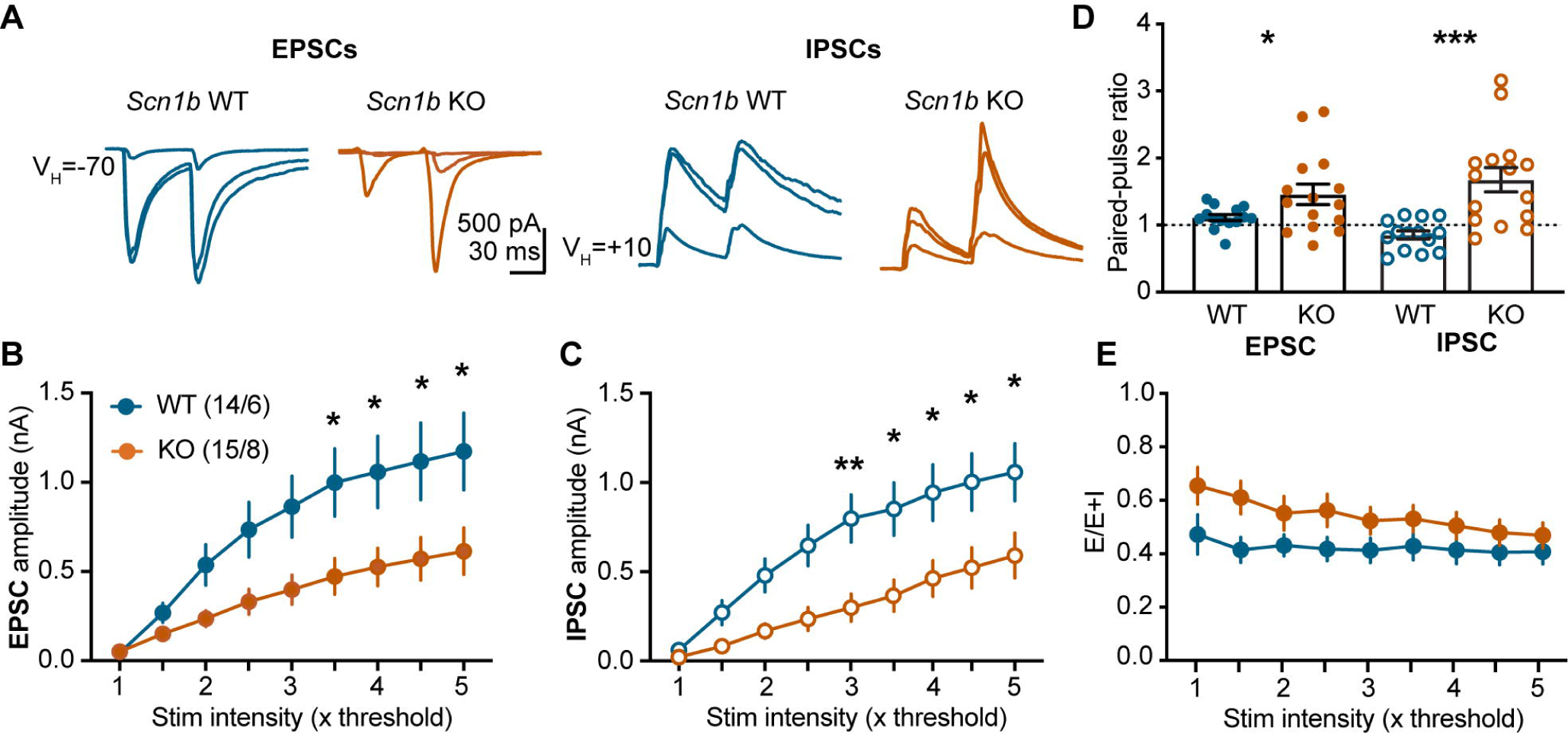
Smaller synaptic currents in *Scn1b* KO pyramidal neurons. **A)** Example whole-cell voltage clamp recording responses to paired Schaffer collateral stimulation (50 ms inter-stimulus interval) at 1, 3 and 5 times (overlayed) the minimal stimulation intensity necessary to evoke a reliable EPSC. EPSCs were recorded by voltage-clamping the cell E_GABA_ (−70 mV) and IPSCs at E_glutamate_ (+10 mV). **B)** Summary data of EPSC input/output curve demonstrates that EPSCs were smaller across stimulation intensities in KO neurons compared to WT. N = cells/mice. **C)** Summary data of IPSC input/output showing that IPSCs were also smaller in KO neurons (*p < 0.05; **p < 0.01, Sidak’s multiple comparisons test). **D)** Increased paired-pulse ratio of both EPSCs (left, closed circles) and IPSCs (right, open circles) in KO neurons at 3x stim intensity. *p < 0.05; ***p < 0.001, Mann-Whitney tests. **E)** Summary data of excitatory to inhibitory ratio, measured as the area of the EPSC/(EPSC+IPSC).

We next performed a similar input/output experiment in current clamp to measure SC driven <u>p</u>ost<u>s</u>ynaptic <u>p</u>otentials (PSPs). We used a single input stimulation paradigm similar to that in Figure 5, which produced smaller synaptic current growth functions in KO neurons, and again normalized stim intensity to the smallest intensity that evoked a reliable postsynaptic potential (∼ 1 mV in amplitude), then increased by multiples of that threshold stimulation. No synaptic blockers were present in order to measure a mixed PSP. KO neurons had larger PSPs (APs excluded/truncated) compared to WT (Figure 6 A-B; main effect of genotype: F_(1, 27)_ = 6.476, p = 0.017; stim intensity: F_(8, 216)_ = 48.380, p < 0.0001; stim intensity x genotype: F_(8, 216)_ = 2.301, p = 0.022). We also found that with even a single stimulation, a proportion of KO cells fire, whereas WT neurons do not. Although, the firing probability was not significantly different between genotypes (Figure 6C; main effect of genotype: F_(1, 27)_ = 2.300, p = 0.141; stim intensity: F_(8, 216)_ = 1.922, p = 0.058; stim intensity x genotype: F_(8, 216)_ = 1.922, p = 0.058).

**Figure 6.**
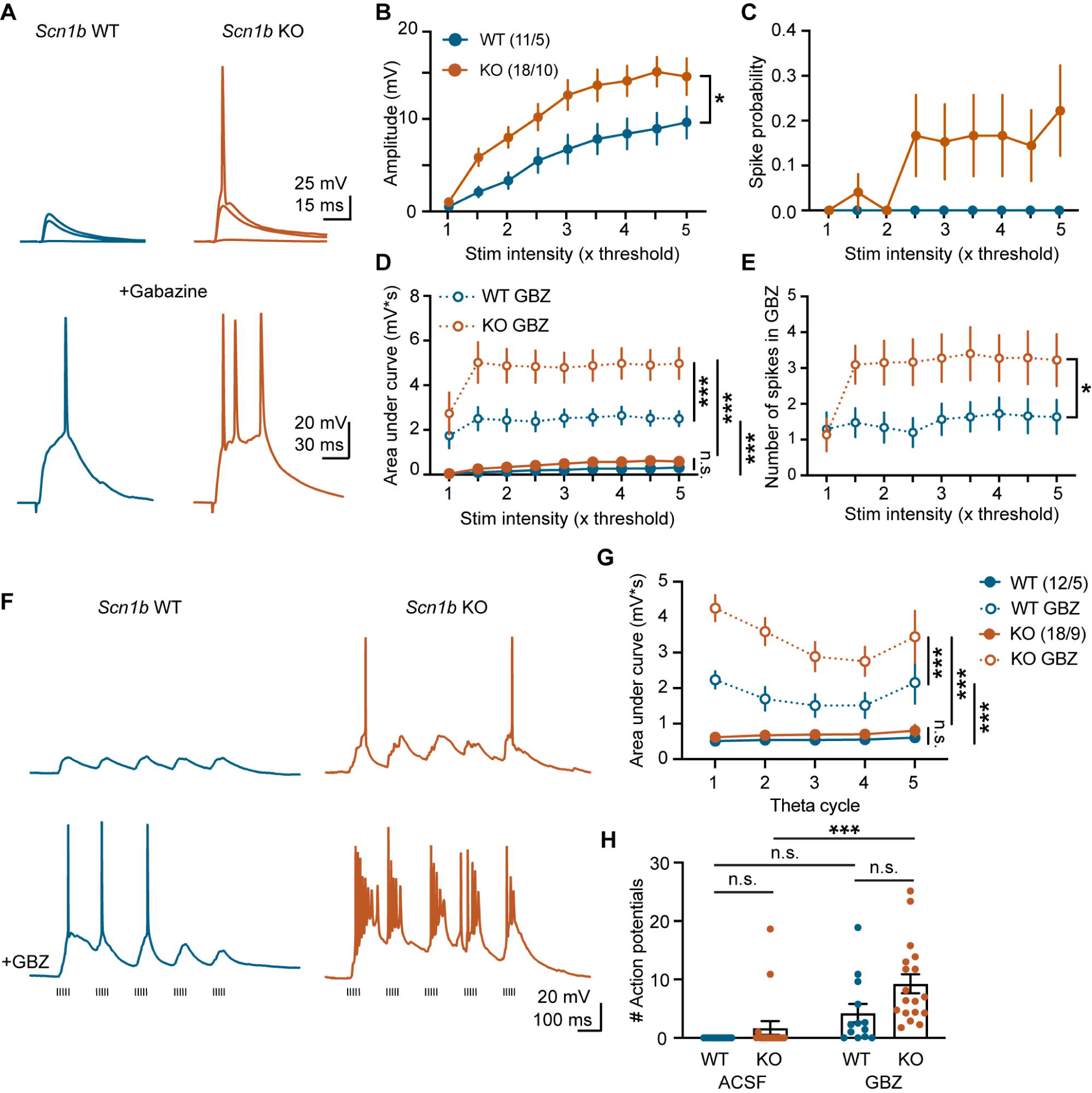
Larger postsynaptic potentials in *Scn1b* KO pyramidal neurons. **A)** Example current clamp recordings of postsynaptic potentials in response to Schaffer collateral stimulation at 1, 3 and 5 times the threshold intensity from WT (left, blue) and KO (right, orange) neurons with GABAergic signaling intact (top). Bottom traces are with GABA_A_ signaling blocked with gabazine (GBZ; 10 μm; 3x threshold stim). **B)** Summary input/output data showing that mixed postsynaptic potentials are larger in KO neurons compared to WT. N = cells/mice. *p = 0.017 for main effect of genotype, two-way RM ANOVA. **C)** Spike probability vs. stim intensity for WT and KO neurons when GABA signaling is intact demonstrating that some KO neurons begin to fire with a single stimulation, and WT neurons do not. **D)** Amplitude of PSPs, measured as area under the curve with spikes truncated. Solid circles are mixed PSPs. Open circles are in presence of gabazine. ***p < 0.001; n.s = not significant (p > 0.05), Tukey’s multiple comparisons tests.**E)** Number of spikes vs. stim intensity in gabazine. *p < 0.05 main effect of genotype, two-way RM ANOVA. **F)** Example traces from *Scn1b* WT (left) or KO (right) neurons in response to theta burst stimulation of Schaffer collaterals (5 pulses at 100 Hz, repeated 5 times, 200 ms inter-burst interval; hash marks below) in standard ACSF (top) or gabazine (10 μm; bottom). **G)** Magnitude of temporally summed PSPs, measured as area under the curve with spikes truncated. Solid circles are mixed PSPs. Open circles are in gabazine. ***p < 0.001; n.s = not significant (p > 0.05), Tukey’s multiple comparisons tests. N = (cells/mice). **H)** Number of action potentials resulting from theta burst stimulation in ACSF and GBZ. ***p < 0.001; n.s = not significant (p > 0.05), Sidak’s multiple comparisons tests.

While our voltage clamp experiments revealed similar excitatory/inhibitory ratios between genotypes (Figure 5E), altered interactions between these inputs could underlie the larger synaptic depolarizations in *Scn1b* KO pyramidal neurons in current clamp. We hypothesized that if the enhanced postsynaptic potentials were due to a reduction in inhibitory control of synaptic integration, blocking inhibition would have a larger effect on WT neurons compared to KOs. We washed on the GABA_A_ antagonist gabazine (10 μM, GBZ; Figure 6A), and found that while responses in both genotypes were increased, gabazine increased the amplitude (Figure 6D) and spiking (Figure 6E) of KO neurons to a greater extent (Figure 6D: main effect of stim intensity: F_(8, 216)_ = 6.777, p < 0.0001; drug: F_(1, 27)_ = 50.10, p < 0.0001; genotype F_(1, 27)_ = 6.469, p = 0.017; stim intensity x drug: F_(8, 216)_ = 3.284, p = 0.002; stim intensity x genotype: F_(8, 216)_ = 1.379, p = 0.207; drug x genotype: F_(1, 27)_ = 4.856, p = 0.036; stim x drug x genotype: F_(8,216)_ = 0.876, p = 0.538; three-way RM ANOVA with post-hoc Tukey’s multiple comparisons tests; Figure 6E: main effect of stim intensity: F_(8, 216)_ = 4.216, p = 0.0001; genotype: F_(1, 27)_ = 3.370, p = 0.077; intensity x genotype: F_(8, 216)_ = 2.568, p = 0.0107; two-way RM ANOVA). This is contrary to our hypothesis of functionally deficient inhibition, suggesting that GABAergic inhibition is controlling a much greater degree of hyperexcitability to synaptic input in the *Scn1b* KO hippocampus. However, this is a single stimulation, and the enhanced synaptic integration seen in Figure 1 could be driven by a breakdown of GABAergic signaling with high frequency stimulation.

To address the role of inhibition in synaptic integration, we repeated the theta burst paradigm in a new set of cells. We again normalized the stimulation intensity such that the first stimulation elicited a consistent PSP of 1-2 mV (as in Figure 1), measured temporal integration of theta burst stimulation in standard ACSF, and then washed on gabazine (10 μM, Figure 6F). We again found that the postsynaptic responses were larger in *Scn1b* KO neurons compared to WT (Figure 6H, main effect of genotype: F_(1,45)_ = 26.59, p < 0.0001, three-way RM ANOVA) and that gabazine increased this response (main effect of drug: F_(1,45)_ = 221.5, p < 0.0001). As with responses to single stimuli, KO neurons showed a much greater increase in activation with inhibition removed than did WT, as revealed by a significant drug x genotype interaction (F_(1,29)_ = 7.945, p = 0.009; full three-way RM ANOVA model: main effect of theta cycle: F_(4, 116)_= 4.774, p = 0.001; theta cycle x genotype: F_(1,116)_ = 0.694, p = 0.598; theta cycle x drug: F_(4, 116)_ = 6.926, p < 0.0001; theta cycle x genotype x drug: F_(1,116)_ = 1.182, p = 0.322; three-way RM ANOVA with post-hoc Tukey’s multiple comparisons tests). Blocking GABA also increased firing in response to theta burst stimulation of SC axons (main effect of drug: F_(1, 58)_ = 19.03, p < 0.0001, two-way ANOVA with post-hoc Sidak’s multiple comparison test), and KO neurons fired more action potentials than WT neurons (main effect of genotype: F_(1, 58)_ = 6.155, p = 0.016), but GABA blockade did not differentially effect one genotype over the other (drug x genotype: F_(1, 58)_ = 1.514, p = 0.224). In summary, both single and patterned stimulation evoked large, long-lasting PSPs and repetitive firing in *Scn1b* KO slices when GABA_A_ signaling was blocked, indicating that GABA effectively dampens runaway excitability in the KO hippocampus. However, because of the recurrent/epileptiform activity in KO slices in gabazine, GABA blockade is not a feasible way to test the role of inhibition in the enhanced synaptic integration see in Figure 1.

Previous studies have reported mixed results on how *Scn1b* deficits alter the excitability of GABAergic interneurons (Reid, Leaw et al. 2014, Hull, O’Malley et al. 2020), and none have examined the recruitment of interneurons during physiologically relevant synaptic stimulation. We focused on two major hippocampal interneuron populations in CA1, parvalbumin (PV+) expressing fast-spiking interneurons that mainly provide feedforward inhibition to the soma or perisomatic regions to regulate pyramidal cell firing, and somatostatin (SST+) expressing interneurons located in stratum oriens, that provide mostly feedback inhibition to the dendrites of pyramidal neurons to regulate synaptic integration (Pelkey, Chittajallu et al. 2017). We crossed the *Scn1b* knockout mouse line with PV-Cre-Ai14-TdTomato and SST-Cre-Ai14-TdTomato reporter mice to identify interneurons in hippocampal slices.

Using a series of current steps, we measured firing properties of CA1 PV+ interneurons (Figure 7A) and found that fast-spiking PV+ neurons from KO slices fired fewer action potentials compared to WT (Figure 7B; main effect of current injection: F_(2.168,90.74)_ = 112.0, p < 0.0001; main effect of genotype: F_(1, 43)_ = 6.384, p = 0.015; current x genotype: F_(7, 693)_ = 4.852, p < 0.0001; two-way RM ANOVA). The mechanism for this change in firing isn’t immediately clear as it was not accompanied by a change in resting membrane potential (Figure 7C; t_(43)_ = 0.744; p = 0.461; t-test), membrane resistance (Figure 7D; t_(43)_ = 0.556 p = 0.581; t-test), voltage threshold (Figure 7E, left; t_(43)_ = 0.873; p = 0.388; t-test), or rheobase (Figure 7E, right; t_(43)_ = 1.211; p = 0.236; t-test). We also found no difference in capacitance of PV+ cells between genotypes (not shown; WT: 134.1 ± 8.890 pF; KO: 134.8 ± 10.91 pF; t_(43)_ = 0.050, p = 0.960). In contrast, firing properties of SST+ interneurons were not different between genotypes (Figure 7 F-G; main effect of current injection: F_(1.60, 81.55)_ = 98.39, p < 0.0001; main effect of genotype: F_(1, 51)_ = 1.968, p = 0.167; current x genotype: F_(12, 612)_ = 0.495, p = 0.918; two-way RM ANOVA). We also found no change in resting membrane potential (Figure 7H; p = 0.663; Mann-Whitney test), membrane resistance (Figure 7I; t_(51)_ = 1.536 p = 0.131; t-test), voltage threshold (Figure 7J, left; t_(51)_ = 0.848; p = 0.400; t-test), or rheobase between genotypes (Figure 7J, right; t_(51)_ = 1.728; p = 0.090; t-test).

**Figure 7.**
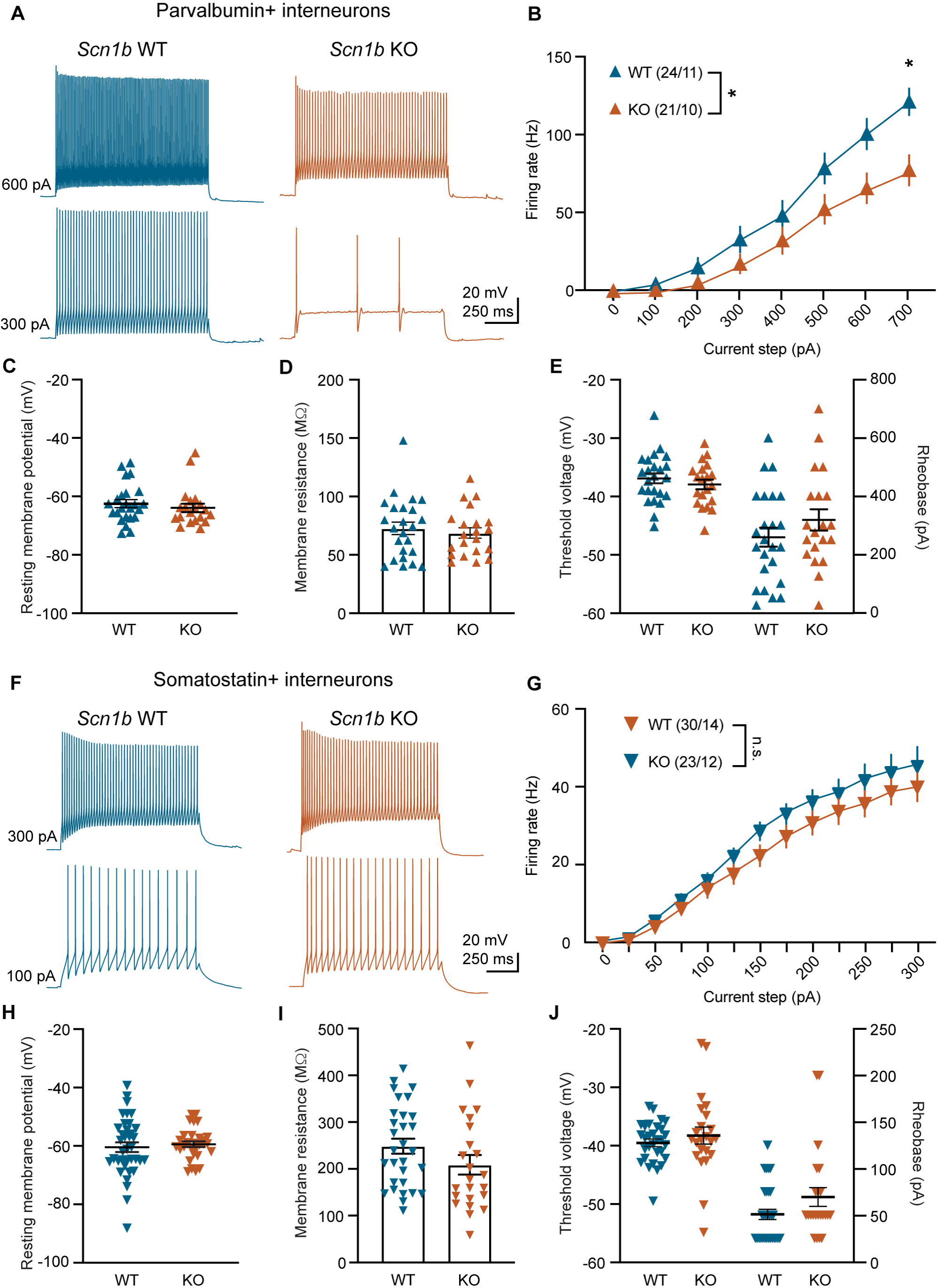
Reduced firing in PV+ but not SST+ interneurons from *Scn1b* KO mice. **A)** Example traces from WT (left, blue) and KO (right, orange) parvalbumin (PV) expressing interneurons in response to current injections of 300 pA (bottom) and 600 pA (top). **B)** F/I plot from PV+ interneurons showing that KO neurons fire less than WT neurons. *p < 0.05, main effect of genotype, two-way RM ANOVA; *p = 0.012 between genotype comparison at 700 pA, Sidak’s multiple comparison’s test. N = cells/mice. **C)** There was no difference in resting membrane potential, **(D)** membrane resistance, **(E)** voltage threshold (left), or rheobase (right) between genotypes in PV+ neurons**. F)** Example traces of from WT (left, blue) and KO (right, orange) somatostatin (SST) expressing interneurons in response to current injections of 100 pA (bottom) and 300 pA (top). **G)** F/I plot from SST+ interneurons. n.s = p > 0.05, main effect of genotype, two-way RM ANOVA. **H)** KO SST+ interneurons showed no difference in **(H)** resting membrane potential, **(I)** membrane resistance, **(J)** voltage threshold (left), or rheobase (right) between genotypes.

We next tested how these interneuron subtypes are recruited during theta-burst stimulation of stratum radiatum. Like CA1 pyramidal neurons, many CA1 PV+ interneurons receive direct excitation from Schaffer collateral axons, as well as feedforward inhibition. In response to theta-burst stimulation, WT and KO PV+ interneurons had similar responses at the minimal stimulation intensity (Figure 8 A-B; main effect theta cycle at 1x stim intensity: F_(4, 110)_ = 0.273, p = 0.89; main effect of genotype: F_(1, 110)_ = 2.001, p = 0.160; theta cycle x genotype: F_(4, 110)_ = 0.192, p = 0.942). As stim intensity was increased as in earlier experiments, PV+ cells from KO mice displayed smaller PSPs (avg AUC at 5x stim: WT = 7514 ± 1112 mv*ms; KO = 4378± 832.2 mv*ms; t_(22)_ = 2.258, p = 0.034, t-test) and fired fewer action potentials than those from WT mice (Figure 8 A-B; main effect theta cycle at 3x stim intensity: F_(4, 110)_ = 0.076, p = 0.989; main effect of genotype: F_(1, 110)_ = 4.682, p = 0.033; theta cycle x genotype: F_(4, 110)_ = 0.033, p = 0.998; main effect theta cycle at 5x stim intensity: F_(4, 110)_ = 0.889, p = 0.473; main effect of genotype: F_(1, 110)_ = 16.18, p = 0.0001; theta cycle x genotype: F_(4, 110)_ = 0.255, p = 0.906; two-way RM ANOVA). These data suggests that PV+ interneurons are less excitable and much less likely to be recruited during physiological activity, and a reduction in fast (peri)somatic inhibition may contribute to the increased integration and firing in KO pyramidal neurons seen in Figure 1.

**Figure 8.**
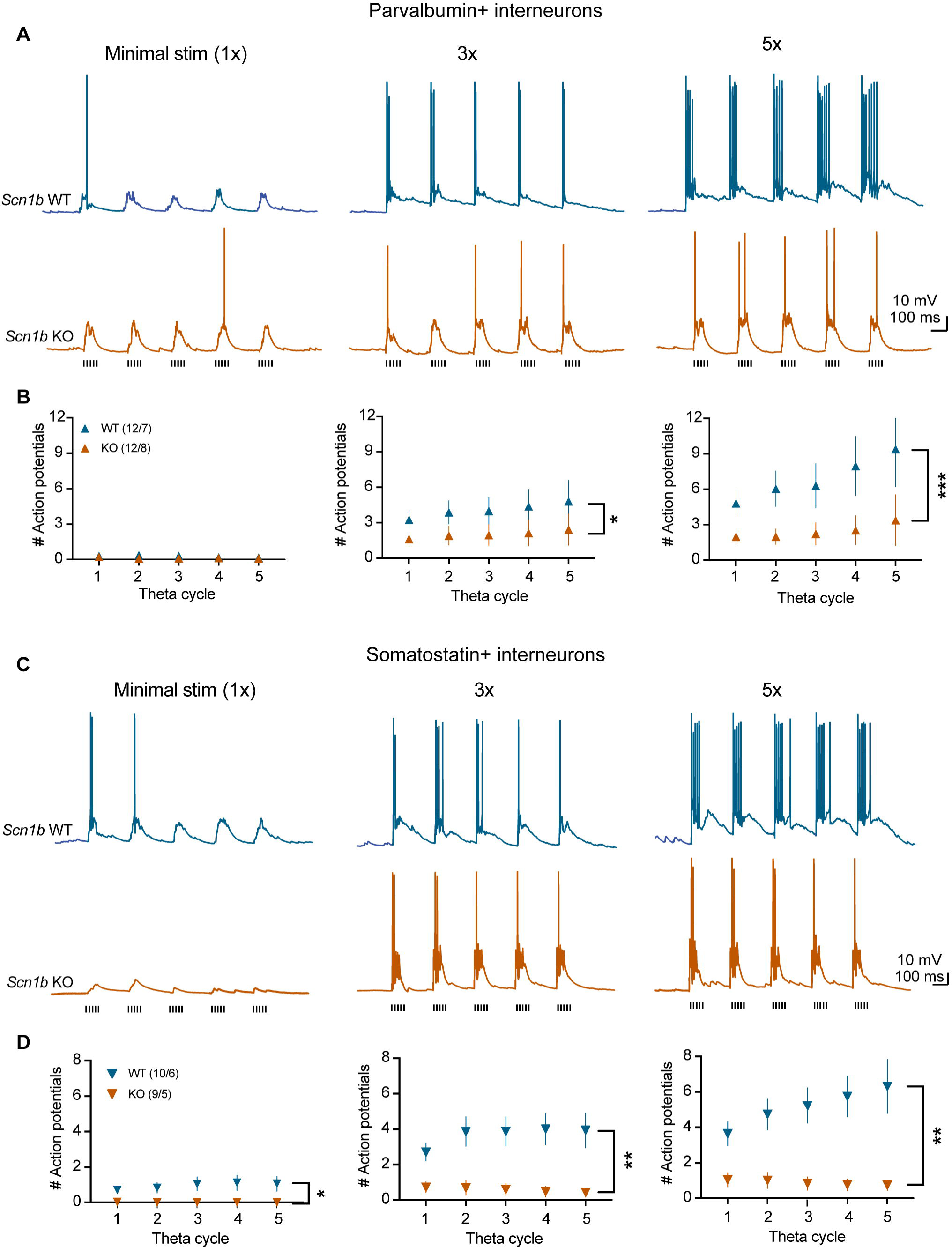
PV+ and SST+ interneurons are less likely to be recruited by theta burst stimulation in the *Scn1b* KO hippocampus. **A)** Example PSPs recorded from PV+ interneurons from WT (top, blue) or KO (bottom, orange) in response to SC theta burst stimulation at the threshold stimulation intensity (1x; left), 3x (middle), and 5x the threshold intensity (right). **B)** Number of action potentials per theta cycle at the 1x (left), 3x (middle), and 5x (right) stim intensities. N = cells/mice. *p < 0.05; ***p < 0.001; main effect of genotype; two-way RM ANOVA. **C)** Example PSPs recorded from SST+ interneurons from WT (top, blue) or KO (bottom, orange) slices in response to SC theta burst stimulation at 1x (left), 3x (middle), and 5x (right) the threshold intensity. **D)** Number of action potentials per theta cycle at the 1x (left), 3x (middle), and 5x (right) stim intensities. N = cells/mice. *p < 0.05; **p < 0.01; main effect of genotype, two-way RM ANOVA.

We also investigated the recruitment of SST+ interneurons by theta-burst stimulation. Because these cells receive excitatory synaptic input from CA1 pyramidal neurons (Müller and Remy, 2014) and their measured intrinsic properties were normal, we hypothesized that the increased firing of KO pyramidal neurons (Figure 1) would increase the recruitment of SST+ interneurons, possibly providing strong inhibitory control even though inhibitory synaptic inputs onto pyramidal neurons were diminished in amplitude. We again performed the theta-burst stimulation paradigm in stratum radiatum while recording from SST-Cre-TdTomato+ neurons in stratum oriens from *Scn1b* WT and KO mice (Figure 8C). Unexpectedly, we found that despite normal intrinsic firing properties, recruitment of SST+ neurons was greatly diminished in KO mice, firing fewer action potentials than those from WT mice across all stimulation intensities (Figure 8A-B; main effect theta cycle at 1x stim intensity: F_(1.1, 18.7)_ = 0.961, p = 0.348; main effect of genotype: F_(1, 17)_ = 7.318, p = 0.015; theta cycle x genotype: F_(4, 68)_ = 1.170, p = 0.332; main effect theta cycle at 3x stim intensity: F_(1.6, 26.7)_ = 2.835, p = 0.087; main effect of genotype: F_(1, 17)_ = 15.20, p = 0.001; theta cycle x genotype: F_(4, 68)_ = 4.794, p = 0.002; main effect theta cycle at 5x stim intensity: F_(1.8, 30.4)_ = 1.601, p = 0.219; main effect of genotype: F_(1, 17)_ = 13.37, p = 0.002; theta cycle x genotype: F_(4, 68)_ = 2.938, p = 0.027; two-way RM ANOVA). Because these cells provide inhibition to the dendrites of CA1 pyramidal neurons to regulate synaptic integration, the lack of recruitment of SST+ neurons may contribute to the enhanced synaptic integration in pyramidal neurons. Overall, these results show that simply recording somatic responses to current steps or measuring synaptic currents does not provide a complete picture and can mask changes in neural processing at the cellular level that give rise to changes in how a neural circuit processes ongoing activity.

## Discussion

Our data reveal that loss of *Scn1b*, a causative gene for severe epileptic encephalopathy disorders, leads to changes in subthreshold and suprathreshold intrinsic properties, excitatory and inhibitory synaptic physiology, and synaptic integration that were not restricted to a single neuron type or consistent across cell types. The result is a dramatic increase in the input/output function of the hippocampus, likely underlying the myriad neurological deficits experienced by people suffering from these disorders, even in interictal periods.

The known biochemistry of β1 provides interesting clues toward, but not a complete explanation for the complex results of our study. While *SCN1B* was identified and named for its interactions with alpha pore-forming subunits of voltage-gated sodium channels (Hartshorne and Catterall, 1984; Messner et al., 1986; Isom et al., 1992), β1 also interacts with and modulates the physiology of potassium channels involved in action potential repolarization and dendritic synaptic integration (Deschênes and Tomaselli, 2002; Marionneau et al., 2012; Nguyen et al., 2012). The extracellular domain of β1 serves as a cell adhesion molecule (Davis et al., 2004; Brackenbury et al., 2010; Patino et al., 2011), and the intracellular domain interacts with fyn kinase to regulate neurite outgrowth (Brackenbury et al., 2008) or can be cleaved and translocate to the nucleus to alter transcription (Haworth et al., 2022). The multiple functions of this protein indicate an array of pathways through which neuronal physiology could be regulated/disrupted, complicating the question of whether these phenotypes are primary to loss of protein function, compensatory mechanisms, or driven by seizures that wrack the circuitry.

For example, findings that are likely a direct result of loss of β1 are reduced action potential amplitude in CA1 pyramidal neurons and decreased firing in PV+ interneurons, consistent with a role for β1 in stabilizing surface expression of and increasing conductance through sodium channels. A less clear finding is the increased sag voltage in *Scn1b* KO pyramidal neurons. There is no known direct interaction between β1 and HCN channels, which mediate I_h_ and voltage sag. HCN channels in pyramidal neurons are known for their gradient of increasing expression from soma to distal dendrites where they suppress synaptic integration and activity (Magee, 1998; Lörincz et al., 2002). The mechanism for upregulation of HCN channels in the somatic compartment described here is not clear, but functionally decreases input resistance in opposition to lower capacitance. It may be that these channels are present in the soma due to disrupted dendritic trafficking, which would amplify synaptic inputs (Magee, 1999; Shin et al., 2008), consistent with the enhanced synaptic integration revealed in our data. Alternatively, upregulation of HCN channels may be a homeostatic response to counteract enhanced excitability (van Welie et al., 2004; Fan et al., 2005), or a developmental delay, as somatic HCN channel density is higher in immature neurons to counteract high input resistance (Yang et al., 2013). Interestingly, voltage sag upregulation was recently reported in the*Scn1a* mouse model of Dravet syndrome (Jones et al., 2022), while loss of HCN1 gene function can cause Dravet syndrome and related epileptic encephalopathies (Nava et al., 2014; Bonzanni et al., 2018).

Previous studies of *Scn1b* loss on neuron physiology reveal mixed results varying by cell type, brain area, preparation, and model. For example, Reid and colleagues (2014) found increased excitability in <u>L</u>ayer (L) 2/3 cortical and subicular pyramidal neurons, but normal physiology in CA1 and L5 cortical pyramidal neurons in a *Scn1b* loss-of-function disease mutation model. *Scn1b* KO L6 and subicular pyramidal cells have also been shown to fire more action potentials with smaller current injections compared to WT, but go into depolarization block and fire less with larger current steps (Hull et al., 2020). Hull and colleagues (2020) found no change in L5 pyramidal cell firing, whereas *Scn1b* KO L5 pyramidal neurons displayed increased firing in response to current injections in another study (Marionneau et al., 2012). In CA1 pyramidal neurons, we found only a modest increase in firing in response to larger amplitude current steps, without a change in resting membrane potential, voltage threshold, or rheobase. These data collectively indicate that subtle increases in intrinsic somatic excitability of pyramidal neurons likely contribute to the epileptic phenotype of *Scn1b* loss but is not the only source of dysfunction.

Another level of neuronal processing altered in *Scn1b* KO hippocampus is synaptic transmission. Our data demonstrate a surprising reduction in postsynaptic current amplitude of both excitatory and inhibitory currents, and these are reduced in a similar manner without changing the excitatory to inhibitory ratio across a broad range of stimulation intensities. While smaller, the currents are substantially more facilitating in KOs compared to WT, likely contributing to the enhanced integration of inputs and indicating changes to the physiology of neurotransmitter release. These smaller synaptic currents combined with the relatively subtle increases in somatic excitability would not be expected to produce the greatly enhanced excitability of KO pyramidal neurons in response to even single synaptic stimulations. This suggests that dendritic physiology and processing of incoming synaptic inputs is altered in *Scn1b* KO pyramidal cells. The intrinsic physiological properties of dendrites are unique and independent from the cell body, as dendritic morphology and distribution of ion channels act to dampen or enhance signal propagation (Spruston, 2008). We did not find differences in dendritic length or gross dendritic structure of *Scn1b* KO pyramidal cells, and while we can’t rule out less obvious changes in dendritic morphology, we hypothesize that changes to dendritic ion channels play a role in the enhanced synaptic integration found here. There are known interactions between β1 and voltage-gated sodium and potassium channels specifically involved in the control of dendritic excitability (Brackenbury and Isom, 2011; Marionneau et al., 2012; Nguyen et al., 2012), and our data provide evidence of altered physiology of another primary controller of dendritic excitability: HCN channels. Our data reveal further qualitative hints of dendritic hyperexcitability: KO neurons exhibit prolonged depolarization (plateaus) and complex spikes in response to high intensity theta burst stimulation or when inhibition is blocked (see Figures 1A and 6F), which suggests enhanced nonlinear dendritic responses (Spruston, 2008). Alterations in dendritic ion channel distribution and excitability will be explored in our future work.

In another monogenic Dravet syndrome model, the *Scn1a* haploinsufficiency model, a widespread reduction in inhibition due to decreased excitability of several classes of inhibitory interneurons is the consensus phenotype (Catterall, 2018), although the underlying phenotypes are more complicated (Favero et al., 2018; Almog et al., 2021; Chancey and Howard, 2022; Jones et al., 2022; Mattis et al., 2022). Likewise, Hull et al. (2020) found that deletion of*Scn1b* specifically from PV+ interneurons was sufficient to cause seizures and 100% mortality in mice, but the limited studies examining loss of β1 on interneuron function found mixed results, complicating our understanding of the underlying physiology. Hull and colleagues (2020) showed a substantial decrease in firing of cortical parvalbumin positive interneurons and decreased inhibitory synaptic transmission onto cortical pyramidal neurons in a conditional *Scn1b* knockout model, while Reid *et al*. (2014) found no change to hippocampal fast-spiking (putative PV+) interneurons in a *Scn1b* mutant model. Here, we show that PV+ fast-spiking interneurons from KOs had decreased firing in response to current injections, while firing properties in SST+ stratum oriens interneurons were normal, indicating that loss ofβ1 has cell specific effects. We measured the recruitment of these interneurons by physiologically relevant theta-patterned synaptic stimulation, and found that recruitment of both subtypes of interneurons is greatly diminished in *Scn1b* KO slices. Despite these changes to interneuron physiology, blockade of GABA_A_ receptors still caused a massive increase in activity in KO pyramidal neurons compared to WT in response to synaptic stimulation. Thus, synaptic inhibition is clearly exerting a great deal of control over the excitability of pyramidal neurons in the KO brain during ongoing activity. Assessing the role of inhibition is further complicated by data suggesting that the developmental shift in chloride reversal potential is delayed in*Scn1b* KOs (Yuan et al., 2019), which is mitigated by our whole cell recordings. Overall, these data provide further evidence of the complex interplay required for information processing within even a small neural circuit. On top of changes to pyramidal neuron intrinsic and synaptic physiology, differential changes to the physiology of interneuron subtypes and recruitment of inhibition are altered, resulting in enhanced synaptic integration and action potential output from CA1.

Neuronal information processing relies on interplay between synaptic properties, dendritic intrinsic properties that amplify or suppress synaptic signals to integrate them over space and time, and firing properties that produce neuronal output. Disrupting this interplay can result in epilepsy and other neurological disorders. We have shown that loss of*Scn1b* results in a drastically inflated input/output relationship, and that there is neuronal dysfunction all levels of neuronal information processing in *Scn1b* KO pyramidal cells: synaptic inputs; inhibitory drive; synaptic integration; somatic intrinsic physiology; and action potential firing. The sum of these changes is a fundamental change in cellular information processing in the hippocampus, likely contributing to the seizures and associated cognitive defects in*SCN1B*-linked epileptic encephalopathies.

## Acknowledgements

This work would not have been possible without the generous gift of the *Scn1b* transgenic mice from Dr. Lori Isom. We would like to thank members of the Howard and Brumback labs for helpful discussions, especially our technicians who aided in animal care: Mendee Geist; Alexandra Munson; Meredith McCarty; Aurora Weiden; Madelynne Campbell; Joy Adler. This work was funded by an American Epilepsy Society (AES) Young Investigator award to MAH, an AES Postdoctoral Fellowship to JHC, a Dravet Syndrome Foundation Postdoctoral Fellowship to JHC, a University of Texas at Austin TIDES Undergraduate Research Fellowship to AAA, and NIH/NINDS NS112500 to MAH.

